# Viability of carcass removal as an option for offsetting the incidental take of golden eagles (*Aquila chrysaetos*) at wind energy facilities

**DOI:** 10.1101/2022.12.21.521393

**Authors:** Eric V. Lonsdorf, James S. Gerber, Deepak Ray, Steven J. Slater, Taber D. Allison

## Abstract

As wind energy expands to achieve the United States’ net zero emission goals, compensatory mitigation will be required to offset negative impacts to birds and bats. The golden eagle (*Aquila chrysaetos*) is particularly susceptible to collision with wind turbines but only one option for offsetting mortalities has been approved by the U.S. Fish and Wildlife Service despite many sources of anthropogenic-caused mortality. Here, we update a previously developed vehicle-collision model with empirical data and integrate a resource equivalency analysis so that removal of road-killed game animals can be used as mitigation to offset incidental mortality. We parameterized the golden eagle behavior parameters using camera-traps placed at roadside carcasses. We quantified the effects of different carcass-removal schemes based on vehicle and carcass characteristics observed for the state of Wyoming. Our model results indicate that while eagles saved per carcass removed depends on removal interval and vehicle traffic volume, carcass removal is a viable mitigation strategy; up to seven eagles could be saved per year in some counties. While some uncertainty remains about the precise credit received from each carcass removed, delaying the inclusion of additional mitigation methods prevents opportunities for conservation action. An adaptive management program could be a way forward where management and monitoring are combined to further improve estimates of mitigation credit.

---

Wind energy has been a growing source of energy in the United States and with the recent passage of the Inflation Reduction Act, will likely continue to expand. A recent report on the potential for the U.S. to meet a net zero carbon emissions goal suggests that the country will need to increase energy production from wind turbines by 2.5 to 3 times the current installed capacity (Larson et al. 2020). Unfortunately, wind turbines can also have negative impacts on wildlife such as birds and bats (Allison et al. 2019), causing mortality, referred to as incidental take. Eagles are particularly susceptible to incidental take, and the U.S. Fish and Wildlife Service (USFWS) has concluded that the U.S. golden eagle (*Aquila chrysaetos*) population is limited by anthropogenic mortality (USFWS 2016): any increase in mortality could lead to a population decline inconsistent with the preservation standard set by the USFWS in the proposed revised Eagle Rule (see USFWS 2022a, 2022b). Thus, any permits to take golden eagles at wind energy facilities must accomplish “no net loss” through implementation of actions that avoid, minimize, and offset the predicted take.

Despite the many sources of anthropogenic-caused mortality for eagles, retrofitting electric power poles to prevent electrocution of eagles is the only current option approved in the proposed Eagle Rule revision for wind projects seeking permits (USFWS 2013). Having just one option can limit the ability of wind energy companies to implement the mitigation options required for compliance with the Rule. Other sources of mortality and reduced reproduction include lead poisoning from scavenged gut-piles, loss of habitat and prey sources, and collisions with vehicles while scavenging on roadkill (Allison et al. 2017), but these mitigation options are rarely utilized because their application lacks guidance from the USFWS.

Although requirements for approval of mitigation options have not been stated formally, we assume the approach used to translate mitigation actions into golden eagle offset credits would benefit from following three interrelated criteria. First, the mitigation actions must be scientifically rigorous and transparent, such that the logic has been subject to review and collectively agreed upon (Allison et al. 2017). Second, the efficacy of a mitigation approach needs to be verified such that the predictions are defensible, that models have been tested against observation data, and that the level of uncertainty is sufficient for use with mitigation (Cochrane et al. 2020). Finally, the impact of mitigation should be measurable in units identical to the predicted take allowing for easier comparisons to show that the mitigation is equivalent to or greater than the take to be offset, i.e., we must simply be able to translate the number and frequency of carcasses removed into golden eagles saved. The Service provides an example in the Eagle Conservation Plan Guidance of how to use Resource Equivalency Analysis (REA) to calculate lost eagle-years due to electrocution mortality (USFWS 2013) and achieve the goal of offset equivalence to take.

Lonsdorf et al. (2018) developed predictive frameworks for predicting mortality due to collisions from vehicles while an eagle is scavenging on roadkill. The structure of the vehicle collision model was vetted by experts and is logical and straightforward as it lays out the steps that could lead to a vehicle collision: given data from departments of transportation on the occurrence of roadkill, the model describes the number of eagles detecting roadkill, their feeding time on the carcass, the number of vehicles near the carcass and then predicts the probability of an eagle being hit. While there is confidence in the structure of the model, few data were available to parameterize the model so experts were surveyed to elicit functional forms and parameter values. The resulting estimates for mortality rate had considerable uncertainty and were not integrated with an REA. The authors concluded that prioritizing research to update the relationship between eagles and carcasses would increase the reliability of predictive modeling efforts and specific mitigation values.

Two recent studies have since been published whose results could be leveraged to refine the vehicle collision model to increase its acceptance and use as an offset option for eagle take. First, Slater et al. (2022) recently completed an assessment of golden eagle behavior at carcasses along roads. Using camera-traps, they recorded eagle use at more than 150 carcasses along roads in Oregon, Wyoming, and Utah. These data could be used to estimate the functional forms and parameter values for eagle-carcass use in the vehicle collision model. Second, Millsap et al. (2022) evaluated survival rates and the cause of death for golden eagles, collating data from over 500 eagles fitted with transmitters. Among the several causes of death from anthropogenic sources, their data suggested that collisions with vehicles lead to a 1% annual mortality rate with most of these deaths occurring in the winter scavenging season. Millsap et al.’s work provides a constraint to the outcomes that can be used to identify plausible parameter values. Together, these two studies could help improve confidence in the parameter estimates and functional forms, providing necessary verification of the model’s predictions.

Here, using insights and data from these two recent studies to improve confidence in the output of the Lonsdorf et al. (2018) vehicle collision model, we describe our efforts to improve the ability to use carcass removal as a mitigation option. We integrate the updated model with the REA to translate the model output into mitigation credit. We develop a workflow based on nationally available traffic data and apply the workflow to a case study in Wyoming to illustrate the model’s potential application. We evaluate the potential effectiveness of roadkill removal to offset incidental take by addressing two related questions: 1) How does the effectiveness of the option vary with the strategy taken (location and amount of effort) and 2) Is the effectiveness enough for roadside carcass removal to be a viable option for mitigating incidental take?

## METHODS

We developed a model (Figure 1) of Golden Eagle (GOEA) mortality due to vehicle collisions. This model is conceptually similar to the model of Lonsdorf et al. (2018) but parameterized not by expert elicitation but rather by a combination of empirical and inferred parameters. The empirical parameters were derived from a large dataset of eagle/carcass roadside interactions obtained by HawkWatch International (HWI) and the inferred parameters are chosen to ensure that GOEA mortality agrees with observations (in Millsap et al. 2022).

**Figure 1.**
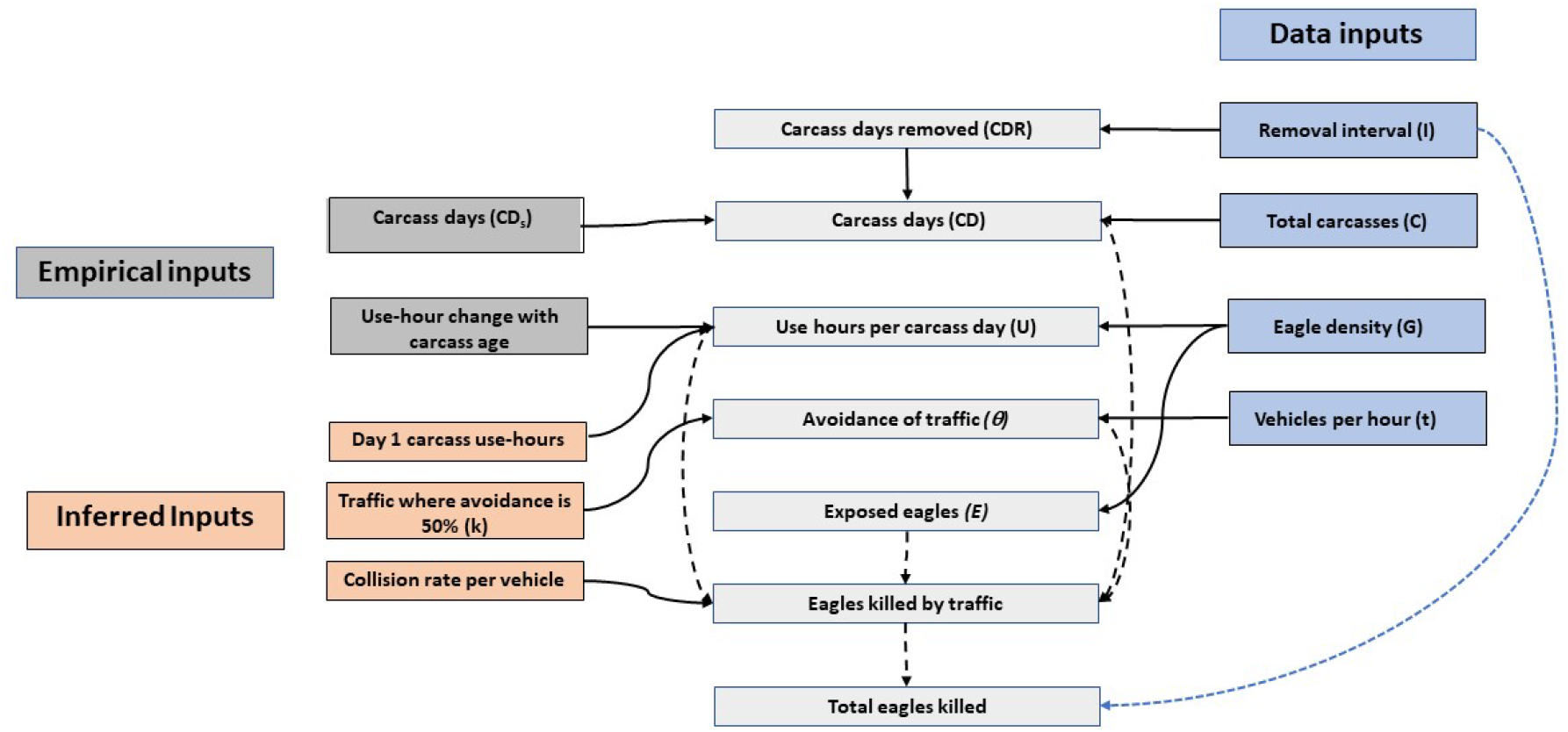
Model diagram illustrating the cause-to-effect relationships (directional arrows) between input and output parameters in the golden eagle vehicle collision model for Wyoming, USA. Blue boxes represent input data, dark grey boxes are parameter estimates derived from empirical studies presented in this paper, salmon-colored boxes are parameter estimates inferred from results from other studies and light grey boxes are model calculations. Solid lines show how parameter values are integrated into equations and dashed lines indicate where the results of one equation are integrated into another. The number of eagles killed is summed over all road segments in a simulation. Adapted from Lonsdorf et al. (2018).

Like the Lonsdorf et al. (2018) model, there are three external factors that set the stage for the model: 1) the number of GOEA at risk which is in turn a function of the density of eagles; 2) the number and size of deer carcasses expected; and 3) the road structure on which the carcasses are found represented by traffic volume *t*. The new model structure diverges from the previous model by simulating the fate of a carcass and eagle behavior each day for a season. Modeling carcasses each day allows us to be explicit about carcass removal strategies and keep track of the number of carcasses collected and the mitigation created by removing each carcass. The simulation models each carcass twice: with and without a given carcass removal strategy.

### Model description

Our updated approach calculates M, the total number of eagles killed, by determining risk created by a single carcass and then summing over the risk of each of N total carcasses. This is expressed by the following equation:

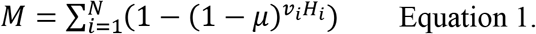

where μ is the per-vehicle-per-eagle collision probability, *v_i_* is the traffic volume in average vehicles per hour (vph) where carcass *i* occurs, and *H*_*i*_ is the total number of eagle use-hours per carcass. This equation is conceptually identical to the power function in equation 1 of Lonsdorf et al. (2018), with the exponent representing the number of vehicles passing scavenging eagles and the base representing the survival probability. However, the nature of the individual terms is different. The total number of eagle use-hours per carcass, *H*_*i*_ is calculated as a sum over the days that the carcass is available, because the use hours per day *h*_*d*_ is a decreasing function of time:

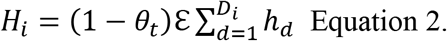

Here, *D*_*i*_ is the number of days carcass *i* is available, *h*_*d*_ is the expected number of hours eagles will scavenge each day carcass *i* is available in the presence of the reference density of eagles, *θ*_*t*_ is the avoidance probability due to traffic volume *t*; and ℇ is a unitless factor modifying the day 1 carcass-use hours. The value of ℇ is constrained by the requirement that modeled overall Golden Eagle mortality due to vehicle strikes agrees with values in the literature (Millsap et al. 2022). The model assumes that road avoidance is a direct and saturating function of average vehicle volume per hour and avoidance affects all eagle age classes equally.

#### Avoidance rate

Eagles can be disturbed by vehicle traffic such that as the volume of cars per hour increases, an increasing portion of the eagles perceive the road as no longer suitable for scavenging. In Lonsdorf et al.’s past work, experts believed that increasing traffic would reduce the number of eagles scavenging and the time spent on the carcass by those eagles that still scavenged. The avoidance probability (**θ*_t_*) for each road with traffic volume *t* is:

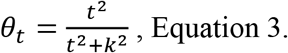

where *t* is the traffic volume (vehicles per hour) and *k* is the traffic avoidance parameter, the traffic volume with 50% traffic avoidance. Experts suggested previously that *k* was between 10 and 35 vehicles per hour (Lonsdorf et al. 2018). Provided that the collision probability *μ* is very small, the factor ℇ can also serve to modify the reference eagle density or *μ* (Supporting Information). Avoidance is 0% with 0 cars/hour and at some traffic volume avoidance approaches 100%.

### Parameter estimates using observed data

We focused our model improvements on updating the representation of golden eagle behavior at carcasses. The HWI data consisted of 1,933 photo interpretations of eagle behavior at roadside deer carcasses, captured with motion-sensitive cameras (see Slater et al. [2022] for details). The interpretations were from a larger set of camera trap observations, and we omitted 12 interpretations due to evident transcription errors. The data included a unique identifier for each camera trap and provided the length of time the camera observed the carcass. For purposes of modeling the camera trap data, we assumed that this corresponded with the actual length of time the carcass was available for golden eagle scavenging.

We recognize that the carcass may have been scavenged or decayed prior to placing the camera (although camera traps were placed preferentially near recent roadkills), and we discuss the potential impacts on the model below. Further, each camera trap was analyzed by a reviewer per frame, and when a golden eagle was seen arriving at the carcass and leaving was noted allowing computation of the time spent scavenging at the carcass. Using the HWI data, our goals are to improve the parameter, *D_i_*, representing the expected number of carcass-days available if there is no removal and to parameterize the number of hours scavenging per carcass day, *h_d_*.

#### Carcass Persistence

When an ungulate is killed by a vehicle collision, the carcass becomes available for scavenging by eagles, but over time, the carcass quality declines as carcass-days increase and eventually the carcass is no longer available or has degraded to the point of having no value to eagles. To estimate the probability that the carcass “persists” we fit a distribution to the HWI data per carcass on the number of observed days they persisted. From the resulting frequency distribution of carcass days from 75 carcasses (Figure 2a), we fit a Cauchy distribution such that the probability that carcass *i* persists *d* days is: 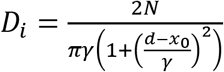, where *d* can take on positive integer values, *γ* is a scaling factor, and *N* is a normalization factor which assures that the discrete sum over probabilities is 1.

**Figure 2.**
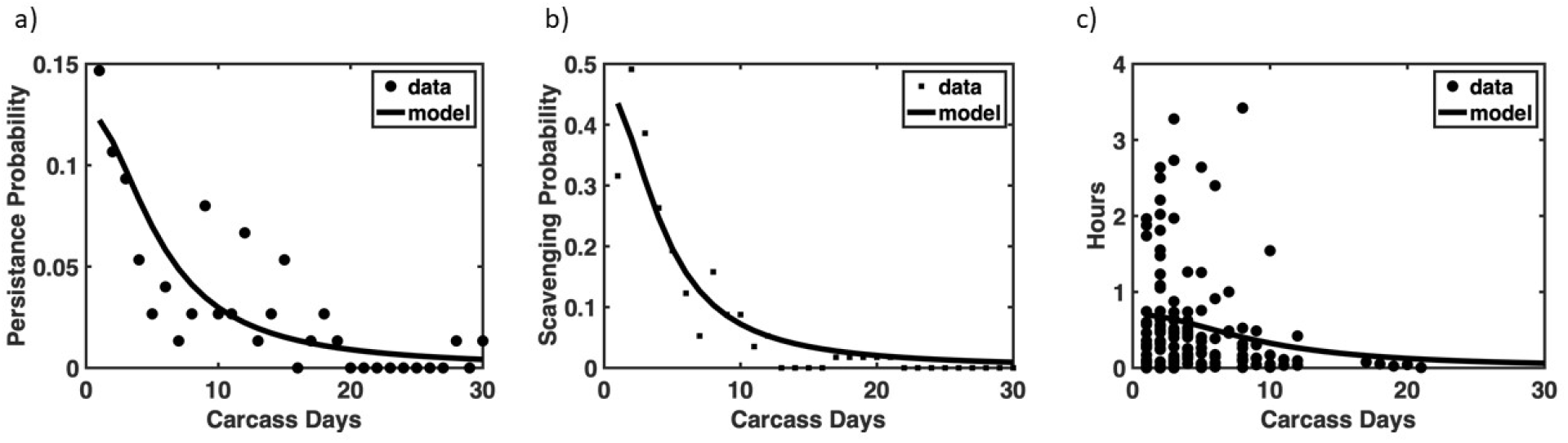
The effect of deer carcass age (days) on persistence, scavenging and use. A. Deer carcass persistence data based on HawkWatch International (HWI) data. B. Daily probability of scavenging. Calculated as the number of carcasses with observed scavenging that many days after camera trap is set divided by the total number of camera traps. Here, the model is a Cauchy distribution with *γ* = 5.56.C. Hours per carcass day if there was scavenging. The model is a Cauchy distribution with parameter *γ* = 4.32.

We chose a Cauchy distribution with offset parameter *x_0_* = 0 because it has the following properties appropriate to the problem at hand:

1. A “fat tail” and thus can represent the data points greater than 30 days.
2. Monotonic decrease from the y-intercept
3. It is a single-parameter distribution, thus decreasing the likelihood of overfitting
4. Lower fitting errors than other distributions meeting the above 3 criteria

We fit by determining the parameter γ which led to the lowest error in an ordinary least squares (OLS) sense, where the OLS calculation was extended out to 100 days as the upper limit for the calculation. We tested several other functional forms for the model (including a Gaussian distribution, a 1/t type of distribution, and an exponential distribution) and confirmed that they led to a higher error or had undesirable properties for carcasses aged beyond a few days (Tables S1-S3, Figs S1-S3, available in Supporting Information).

#### Carcass use-hours

Our assessment of the HWI data found that use-hours can be modeled by a two-step process which retains use-hours’ dependence on carcass age. The first step is to determine the probability that the carcass is scavenged at all; the second is to determine the amount of time spent per day of scavenging when observed. The use-hours per carcass day is the product of these processes, and for simulation purposes we assume scavenging every day with the number of use-hours discounted by the probability of scavenging.

##### Evaluating the probability that a carcass is scavenged

To estimate the probability that a deer carcass is scavenged, we fit a Cauchy distribution to the HWI data in a manner analogous to fitting carcass-days (and choosing a Cauchy distribution for similar reasons, see Supporting Information). Here, for each deer carcass we assessed the number of days (from initiation of photography) where at least one complete scavenging interval was observed.

##### Modeling the use hours per scavenged carcass

To model the expected use hours per scavenged carcass, we fit a Cauchy distribution to the average observed use hours per scavenged carcass as a function of the day (figure 2c). The parameters of that distribution (γ = 9.267) are determined from the HWI data and do not change in modeling vehicle strike. Because there is large variability in use-hours, our use-hour model samples a value from a half-normal (positive values only) distribution with mean value equal to the average use-hours per day.

When applying the use-hours per day function in the model, we assume the decrease in use-hours per day follows the fitted model, but we multiply the entire function by a constant corresponding to a correction for day-1 use hours. In other words, we fit use-hours per day as shown in Figure 3, but multiply by a constant, ℇ, when applying to the model. There are several reasons why such a correction may be necessary, but primarily, the observed use-hours data is of eagles in the presence of traffic, but the model in equation 1 handles decreases in scavenging due to traffic through the avoidance function θ. Other possible reasons include discrepancies in the density of eagles or the effects of carcass aging prior to the start of observations.

**Figure 3.**
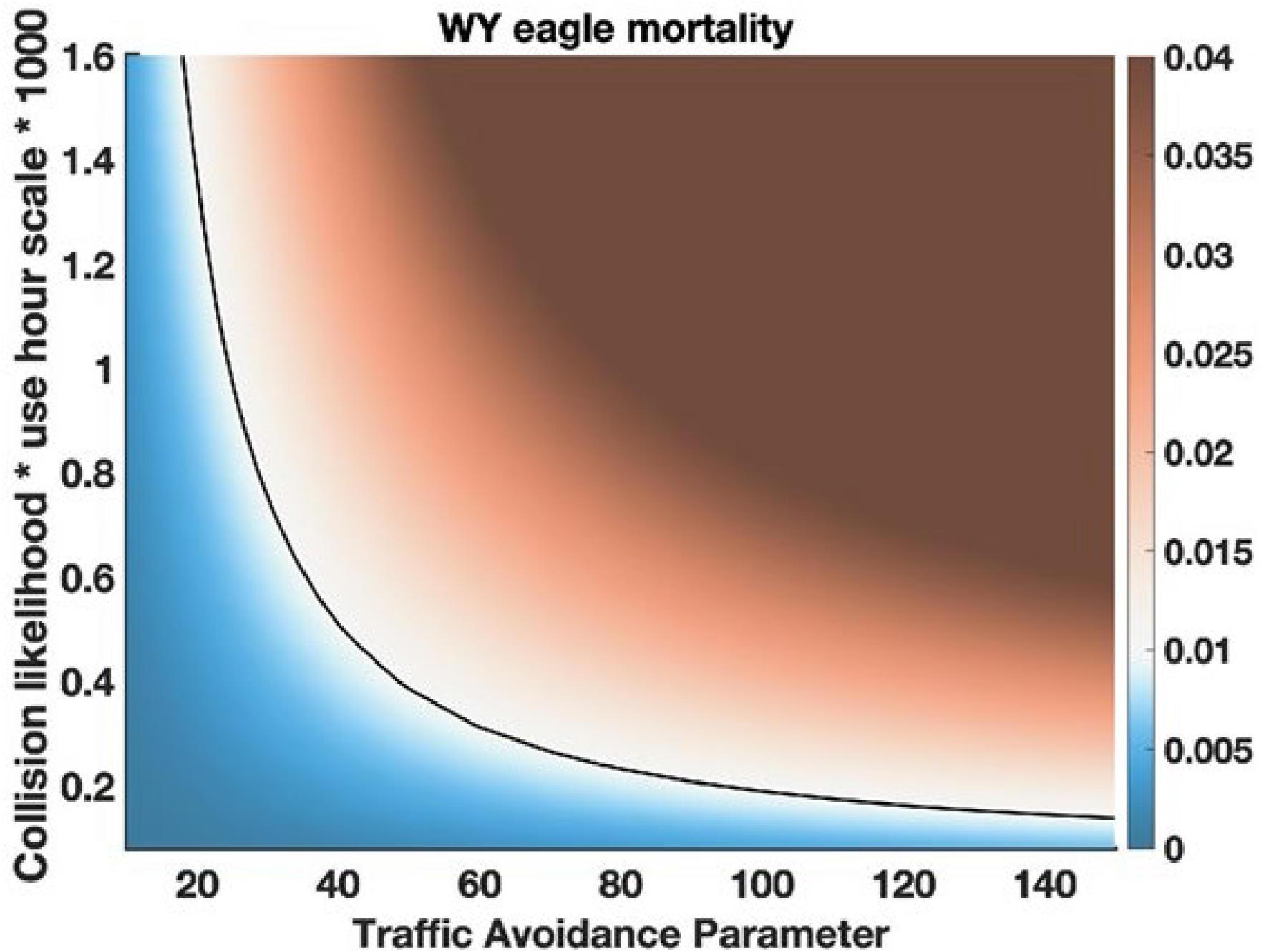
Surface plot of simulated eagle mortality with no carcass removal for a variety of parameters. Traffic avoidance parameter *k* (see Equation 3) varies from 10 to 150. The product of collision likelihood and use-hour scalar varied from 0.0001 to 0.0016. A contour indicating 1% mortality is indicated with a solid black line.

### Simulation model

We implemented a numerical simulation using the Matlab coding language (Mathworks, R2021a). Following Lonsdorf et al. (2018), the simulation leverages vehicle traffic and carcass data from the Wyoming Department of Transportation (WYDOT 2013) and as a first step within each county places a county-specific number of carcasses on roadways with vehicle per hour data characteristics. If the number of carcasses is fractional, there is stochastic rounding to a bounding integer, e.g., an average of 54.3 carcasses has a 70% chance of being simulated as 54 carcasses and a 30% chance of being simulated as 55 carcasses. The distributions of carcass persistence days and use-hours are each sampled, and the total mortality is calculated from equations 1 and 2, first with 0 removals, then with carcass removals at tested intervals: 1, 3, 7, 14 and 30-day intervals). Removal intervals begin randomly relative to carcass days. If the removal day occurs on the final day of carcass availability, this is counted as a carcass removal and the number of use-hours on that final day is reduced by 50%. This procedure is repeated for 500 iterations per interval to allow calculations of 20^th^, 50^th^, and 80^th^ percentile values.

We then used the updated simulation model to evaluate removing carcasses from roads in Wyoming on GOEA vehicle collisions, using the same data from Lonsdorf et al. (2018) which analyzed data from a 10-year study of roadkill on Wyoming highways. We assessed the effect of removal rate from 1 to 30-day intervals applied to all roads within each of Wyoming’s counties.

### Inferred parameters

There are three parameters in the model of Equation 1 (collision probability *μ*, traffic avoidance half-saturation constant k, and day 1 use-hour scaling constant ℇ; also see Figure 1). Provided that eagle-vehicle collisions are rare (*μ*<<1) the parameters *μ* and ℇ are effectively linked (see Supplemental Material) and can be treated as a single parameter, ℇ*μ*. We attempted to determine k empirically, but there were not enough data to estimate a distribution or relationship between traffic volume and flushing rates.

There are *a priori*, a range of possible values of *μ* and ℇ for which the formalism of the present model can be used. However, these values can be constrained with knowledge of total expected adult eagle mortality due to vehicle collisions. Millsap et al. (2022) found that half of collision mortalities, where the cause of collision was determined, were from vehicles. Out of 27,281 eagles observed at the start of a year (the 95% credible interval was 23,374 to 31,779), Millsap et al. (2022) estimated 280 would have died each year from vehicle collisions (95% credible interval was 161 to 439) such that mortality rate would be 1.03% (95% credible interval was 0.69% to 1.38%).

We ran the model simulation over a range of values of the traffic avoidance parameter *k*, and the product of the collision probability *μ* and the use-hour scaling constant ℇ. We varied *k* from 10 to 150, expanding the range from expert elicitation, and we ranged the product of ℇ*μ over values that led to the anticipated eagle mortality of 1%.

### Integrating model with Resource Equivalency Analysis

In the Eagle Conservation Plan Guidance (USFWS 2013), the Service provides a resource equivalency analysis (REA) example to calculate compensatory mitigation designed to offset GOEA take at wind energy facilities (https://www.fws.gov/media/golden-eagle-mortality-resource-equivalency-analysis). Using knowledge of GOEA life history, the REA helps to determine mitigation effort required to save a given number of GOEA adults as a function of expected incidental take. For example, Lehman et al. (2010) estimated that electrocution from power poles led to mortality of at least 0.0036 eagles per pole per year. The REA determines the expected net present value of each lost adult eagle which is a function of its lifetime reproductive value. Specifically, the REA uses the expected age-structure and life history to translate the change in per year mortality into direct and indirect impacts of either the loss of a bird or saving a bird, accounting for age-specific reproduction and mortality. Then, using information on the impact of mitigation, it uses the same logic to determine the effort required to offset the expected take. Thus, given the number of eagles saved per retrofitted power pole, the REA can help determine how many total poles need to be retrofitted given the expected take. We applied this framework for carcass removal to estimate the golden eagles saved for each carcass removed. *Modeling mitigation*: To connect to the REA, we sum up the total carcasses removed for each simulation, determine the change in eagle mortality and simply divide the total eagles saved by the carcasses removed to estimate eagles saved per carcass. We then use the REA to determine the expected total benefits of removing a single carcass, compared to the expected REA-estimated total impact of incidental take. The ratio of loss to the total benefits per-carcass removed provides the number of carcasses needed to be removed each year. If the total carcasses removed each year is greater than those expected from observed data, the duration (number of years) of mitigation effort should increase until the total impact of mitigation reaches the target.

## RESULTS

### Carcass persistence and use

We identified 75 carcasses, of which 57 had at least one complete observation of GOEA scavenging. A histogram of days of carcass availability (Figure 2a) was fit to a Cauchy distribution, resulting in a probability of carcass persistence. Golden eagles spent an average 2.4 days scavenging per carcass (total 134 scavenge-day observations). The histogram of these 134 points (normalized by N=57 carcasses) was fit to a Cauchy distribution (Figure 2b). We found that during the first day, nearly 60% of carcasses were scavenged but that probability declined as the carcass ages (Figure 2c). In contrast to the original representation of scavenging, we found that carcass use hours depended on the amount of time the carcass was available such that as the carcass days increased, the maximum use-hours per carcass day tended to decline. For example, if a carcass was available for 1 or 2 days, as several were, HWI’s data indicate that some carcasses had few use-hours but some had higher use. The upper bound of use-hours declined as carcass days increased. Analysis of HWI’s data suggest that two processes can be modeled to represent carcass use as a function of carcass age. First, we can represent the likelihood, which decreases with carcass age, that the carcass is scavenged at all. Second, the length of time an eagle spends on the carcass if there is scavenging also declines with carcass age.

### Comparison to expert elicitation

Originally, the Lonsdorf et al. (2018) model assumes that as the average number of eagles per available carcass increases, the average amount of scavenging time by individual eagles declines somewhat because of competition. The net result is that use-hours per carcass-day gradually increases with increasing eagle density. The observed data showed more carcass days relative to expert elicitation, but fewer scavenging hours per day.

Experts believed that as eagle density increased the use-hours per eagle would decrease. Lonsdorf et al. (2018) represented this assumption with a power function: *U* = *c* × *G ^z^*, where U is use-hours, *c* is a scalar, *G* is the average density of eagles (number/km^2^) in the county, and the power-function scalar, *z*, is set to 0.5 to approximate the decreasing use with increasing density. This function implicitly assumed that the use hours per carcass-day was independent of carcass-days, i.e., the age of the carcass. The golden eagle density in Wyoming is 0.03 per km^2^, and experts suggested that the parameter *c* was between 3 and 15, leading to an estimated range of carcass use-hours per day of 0.5 to 2.6.

### Inferred mortality parameter families

The resulting mortality for this range of parameters (assuming no removals), along with a contour indicating the family of parameter values of k and ℇ*μ* leading to 1% mortality due to vehicle collisions, is shown in Figure 3. The shape of this equal-mortality contour reflects that expected mortality can be obtained in the model with high traffic avoidance and high collision likelihood (small k and large ℇ*μ*) or low traffic avoidance and low collision likelihood (large k and small ℇ*μ*).

### Effect of location and removal interval

The potential mitigation credit gained from carcass removals is strongly influenced by the location of mitigation and the removal interval (Figure 4). Carcasses are not evenly distributed across Wyoming counties and the traffic volume upon which carcasses are found varies among and within counties, too. Thus, the expected maximum number of carcasses that could be removed per year in each Wyoming county varies widely and depends on the removal interval; total carcasses removed with daily removal range from over 250 in Lincoln County to fewer than 10 in Crook County with the median removed in Wyoming of around 60 carcasses in Platte County (Figure 4a). Increasing removal leads to reduced eagle death but again varies across counties with expected annual mortality dropping from ten to less than four eagles in Lincoln with daily removals, but little impact on mortality in Crook County (Figure 4b). Across all Wyoming counties, credit per carcass removed can vary nearly six-fold with a maximum median number of golden eagles saved per carcass removed of around 0.03 for Carbon County and a minimum of 0.005 in Teton County with daily removal (Figure 4c).

**Figure 4.**
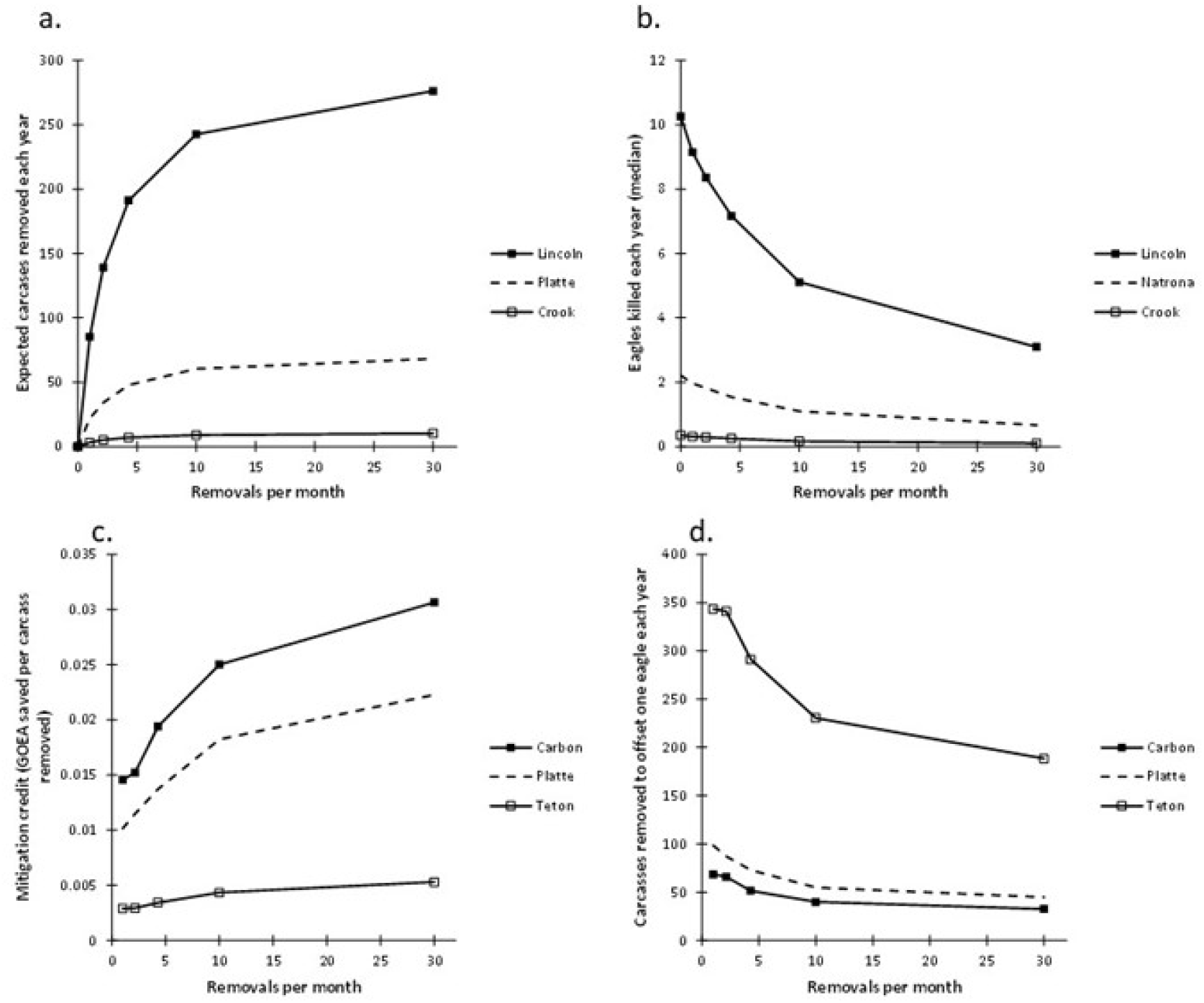
The effect of carcass location and removal interval on range of mitigation impacts. In each panel, the county with maximum mitigation impact is shown with solid line and filled, square marker, the county with minimum impact is shown with solid line and open, square marker and the median county is shown with dashed line and no marker. A. The number of carcasses removed each year, B. The expected eagles killed each year from vehicle collisions, C. The credit expected per carcass removed, and D. the number of carcasses needed to be removed to offset one eagle “taken” each year.

As the interval between carcass removals increases, the number of carcasses removed decreases, the number of eagles saved decreases and the credit per carcass removed decreases. Because we assume that the attractiveness of carcasses declines with increasing age of the carcass, eagles spend most of their total scavenging time in the first few days carcasses are available. As credit per carcass removed declines with fewer removals, it follows that the integration with the REA would indicate that more carcasses will have to be removed to achieve a targeted credit with a maximum of 200 carcasses needed in Teton County with daily removal versus less than 50 in Carbon County (Figure 4d). The difference across counties arises from the expected traffic upon which carcasses are found. Credit is highest on counties with carcasses found on roads with intermediate to low traffic volume (Figure 5).

**Figure 5.**
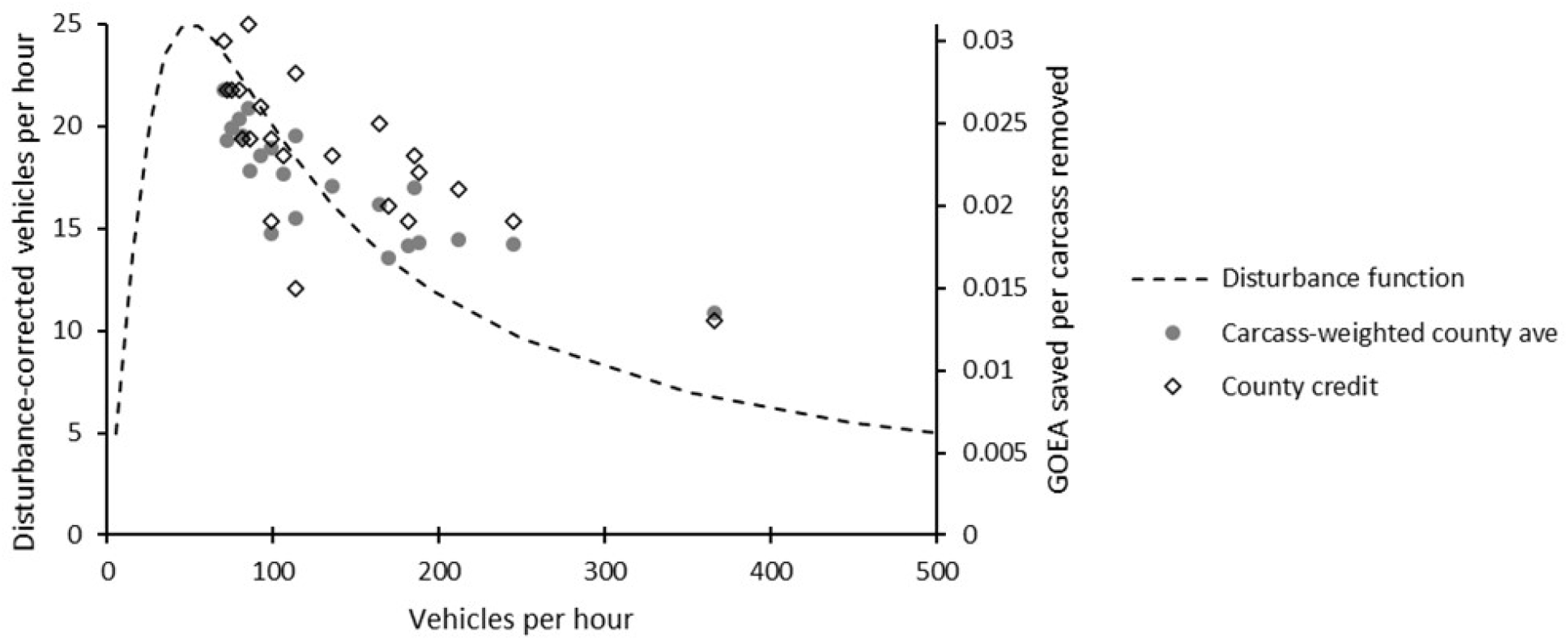
Relationship between county average traffic volume and mitigation credit. As traffic volume increases, the number of cars passing by a carcass along the road increases but will also disturb any eagles feeding on the carcass. The maximum risk of eagle-vehicle collisions, as well as opportunities for mitigation credit per carcass removed, should occur at an intermediate traffic volume.

### Viability for offsetting take

Given that the distribution of deer carcasses and mortality from eagle-vehicle collisions across counties is not uniform, not all counties have equal potential for mitigating incidental take when these results are integrated into the REA (Figure 5, 6). Counties that have both high credit per carcass removed and many carcasses available to be removed have the greatest capacity to provide mitigation credit. Based on the median estimate of credit per carcass and daily effort to remove carcasses, there are several counties where this effort would not be expected to offset the take of a single eagle (figure 6D; white). Other counties, however, could provide annual credits to offset up to 7 eagles killed each year (figure 6D; black).

**Figure 6.**
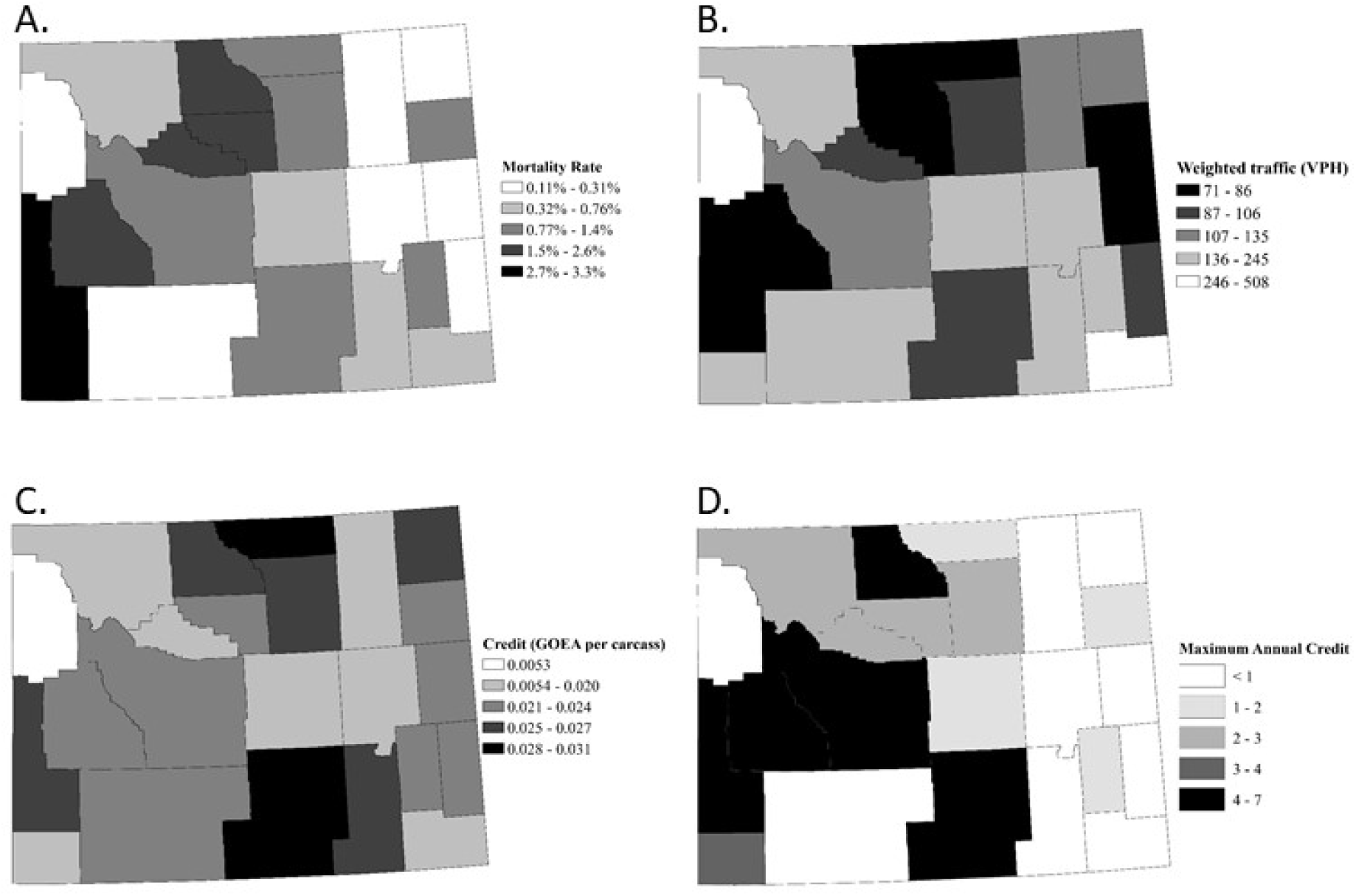
Baseline mortality from golden eagle-vehicle collisions and potential mitigation credit from carcass removals in Wyoming counties. A) While overall mortality in Wyoming is expected to be around 1%, our results indicate that mortality varies across the state with some counties having 3% and some as low as 0.1%. B) The traffic volume where deer carcasses occur also varies across the state. C) The potential mitigation credit per carcass removed varies among counties. D) The total potential credit gained from removing carcasses varies as a function of the credit per carcass removed and the number of carcasses available in the county, with a high of 7 eagles saved per year if all deer carcasses were removed in the county.

## DISCUSSION

With updated parameter estimates, our work shows that carcass removal can be a viable option for mitigation efforts designed to offset incidental take of golden eagles from wind turbines. With empirical observations of mortality from vehicle collisions and behavioral studies of eagle foraging on carcasses, the updated model’s predictions are defensible and the integration of the model results into a USFWS-developed resource equivalency analysis provides clear guidance on effort needed to offset mitigation.

The results also suggest that the per unit effectiveness of carcass removal reducing eagle mortality and the expected number of carcasses available make carcass removal a viable mitigation option. Given the expected 1% golden eagle mortality rate from vehicle collisions, our results suggest that each carcass removed could save up to 0.03 eagles, and thus removing ~34 carcasses with this expected credit per carcass could offset a single eagle taken per year. Past roadkill surveys indicate that there should be sufficient carcasses to remove each year to achieve this level of mitigation. In fact, the credit per carcass removed is nearly equivalent to the effectiveness of a single power pole retrofit despite the fact that a power pole accumulates credit over many years whereas the removal of a single carcass offsets take for only that year. Overall, our results suggest roadkill removal can be adopted as another option for mitigation.

Unlike power pole retrofitting which provides a long-term reduction in risk, carcass removal reduces mortality risk associated with each removal and the risk is likely to return without sustained interventions. Moreover, integrating behavioral studies with the REA indicate that the effectiveness of carcass removal depends on the expected age of the carcass when removed and its location. Because carcasses age, becoming less attractive and less used over time, the expected credit per carcass removed declines as days between removals increase (Figure X). Because increasing traffic can reduce time spent feeding at a carcass, roads of intermediate traffic volume are likely to have higher credit than those with heavy or little traffic, where there is less risk of collision with vehicles. This contrasts with power-poles where eagle density is really the only factor that would affect credit because all poles have assumed constant risk.

There are several reasons to suggest that there should be ample opportunity to use carcass removal as a mitigation. First, traffic data for most major roads are widely available so integrating these data into an analysis like this one should be straightforward. Second, there is increasing understanding of factors that would predict areas where wildlife-vehicle collisions are most likely to occur. Several recent studies indicate that traffic volume and ungulate density are both strong predictors of ungulate collision risk (e.g., Clevenger et al. 2015, Nelli et al. 2018, Mayer et al. 2021), in addition to land cover and seasonality. Deer-vehicle collisions are estimated to cause over $3 billion in damage within the United States (Gilbert et al. 2017) and the ability to predict collision risk should continue to improve. When these locations are identified, our results suggest that there should be sufficient opportunity to use this technique.

While there should be ample opportunities to accumulate mitigation credit, we anticipate that developing and certifying specific mitigation strategies will be potentially challenging. Because carcasses age, there is potential to overestimate the credit if all carcasses removed are credited similarly, regardless of the carcass condition or age. Our results summarize county-wide estimates for credit, but ultimately removal effort is likely to involve driving specific roads regardless of political boundaries. Our results indicate that the strongest predictor of credit is the type of road, so it is possible to use the approach we’ve laid out to provide more refined credit estimates based on road traffic volume. Since the effort and results of the work will need to be monitored and certified, identifying hot spots of opportunity could be helpful in reducing the scale of effort required to monitor and certify mitigation. For example, a recent assessment by the Wyoming Department of Transportation identified 22 one-mile road segments where more than 10 wildlife vehicle collisions occurred each year (Riginos et al. 2016) with more than 65 ungulate carcasses occurring from the top four, one-mile segments. Depending on the traffic volume of these hot spots, our analysis suggests that up to two eagles could be saved per year from ungulate carcass removals during migration season when most collisions occur, thereby making both the mitigation effort and ability to monitor it easier.

Our analysis indicates that some uncertainty remains about the per carcass credit due to the stochastic nature of the deer-vehicle collision process and parametric uncertainty. We used Millsap et al.’s (2022) estimates for eagle mortality from vehicle collisions and the variation in mortality estimate to parameterize our model. The credit per carcass removed and the effort needed to offset take is highly sensitive to this estimate. For example, if the expected mortality from vehicle collisions is 2%, double the current estimate, the credit per carcass removed would also double. We are reassured that a recent 2-year study by HWI in western Wyoming is in general agreement with the Millsap et al. (2022) eagle mortality estimate (HWI, unpublished data). Another potential source of uncertainty is the dependence of results on the vehicle-avoidance parameter *k*, which translates traffic volume into time spent scavenging on a roadside carcass. Application of these results in areas with traffic density characteristics unlike those of our study area would benefit from further study of the relationship between traffic volume and scavenging behavior (Equation 3).

Finally, our work suggests that meeting the required standards for mitigation may be preventing opportunities for conservation actions. While there is still some uncertainty, one must recognize that the uncertainty distribution of credit is one-sided, i.e., we know that carcass removal could be beneficial, but incentives are delayed because of the standards required to determine more precisely how much benefit. Going forward we recommend an adaptive management approach (Williams 2011, Moore et al. 2011) where *a priori* predictions of carcasses removed and GOEA mortality are compared with observations as mitigation proceeds. This would allow for conservation action and research to be done simultaneously, and credit could be updated as the efforts proceed. Doing so would allow progress toward net zero climate emissions to move forward more quickly while we learn about how best to conserve biodiversity.

## MANAGEMENT IMPLICATIONS

Given recent empirical findings and the results of our updated modeling, carcass removal should be added as an acceptable mitigation option. While the mitigation is called “removal”, the carcasses simply need to be moved away from the roadsides; HWI found that moving carcasses twelve meters from the road allows eagles to continue to scavenge and reduces collision risk (Slater et al. 2022). The data required to determine credit - traffic volume, expected “hot-spots” of roadkill and eagle density estimates - are widely available beyond the example we used in Wyoming. If incentivized, carcass removal banks could be created by organizations that could standardize removal intervals and develop a carcass-quality certification process to make these easier for energy companies. If such a program were created, BACI-style research could be done to monitor the impact of reducing carcasses, and when paired with the modeling work, could be used as adaptive management to address the remaining uncertainty in the parameters.

## ACKNOWLEDGMENTS

Financial support for this research was provided by the National Fish and Wildlife Foundation’s Wyoming Golden Eagle Fund. We thank the Partners and Friends of the Renewable Energy Wildlife Institute for their support of eagle mitigation research.

## CONFLICTS OF INTEREST

The authors declare no conflicts of interest.

## ETHICS STATEMENT

No live animals were harmed or handled as part of this research, and all observations of live eagles were collected passively by cameras or from moving vehicles on established roadways.

## DATA AVAILABILITY STATEMENT

Data available on request from the authors.

## Notes

### Competing Interest Statement

The authors have declared no competing interest.

