## Supporting Information for "Viability of carcass removal as an option for offsetting the incidental take of golden eagles (*Aquila chrysaetos*) at wind energy facilities"

December 13, 2022

### Supporting Information

#### Reduction from 3- to 2- parameter model

Equation 1-3 define a 3-parameter model, with parameters being $\mu, ℇ$and $k$

$M=\sum_{i=1}^{N} \left( 1-\left( 1-\mu\right)^{v_{i}H_{i}} \right)$	Equation 1.

$H_{i}=\left( 1-\theta_{t} \right)ℇ\sum_{d=1}^{D_{i}} h_{d}$ Equation 2.

$\theta_{t}=\frac{t^{2}}{t^{2}+k^{2}}$ Equation 3

Equation 1 can be rewritten using a Taylor expansion as follows:

$M=\sum_{i=1}^{N} \left( \mu v_{i}H_{i}+O\left[ \left( \mu v_{i}H_{i} \right)^{2} \right] \right)$ where $O\left[ \left( \epsilon\right)^{2} \right]$ represents terms on the scale of $\left( \epsilon\right)^{2}$and smaller. Since total mortality is expected to be approximately 1%, these terms are necessarily small and hence $\left( \epsilon\right)^{2}$ will be small, hence it is appropriate to treat $\muℇ$ as a single parameter for purposes of mapping out parameter space. In the numerical simulation, the full form of the model in equation 1 is retained for accuracy.

#### Alternative functional forms

We have modeled the Cauchy distribution $F\sim\frac{1}{\left( 1+\left( \frac{d}{a} \right)^{2} \right)}$, an exponential distribution
$F\sim e^{-ad}$, a Gaussian distribution $F\sim e^{-ad^{2}}$ and a 1/t distribution $F\sim\frac{a}{d}$.  We chose the Cauchy distribution in all cases because it has the low relative error, (calculated with an ordinary least squares (OLS) error calculation with an upper limit of d=100 days) and the desired long-tail behavior. In other words, the Cauchy didn’t have the lowest OLS error in all cases, but did have more plausible representation of the likelihood of scavenging for carcasses older than one week.

Table S1 Ordinary Least Squares error of fit of various functions for distribution of carcass days.

|  |  | Total OLS error |
| --- | --- | --- |
| Exponential distribution | $F\sim e^{-ad}$ | 1.004 |
| Gaussian distribution | $F\sim e^{-ad^{2}}$ | 1.404 |
| “1/t” distribution | $F\sim\frac{a}{d}$ | 1.068 |
| Cauchy distribution | $F\sim\frac{1}{\left( 1+\left( \frac{d}{a} \right)^{2} \right)}$ | 1.000 |

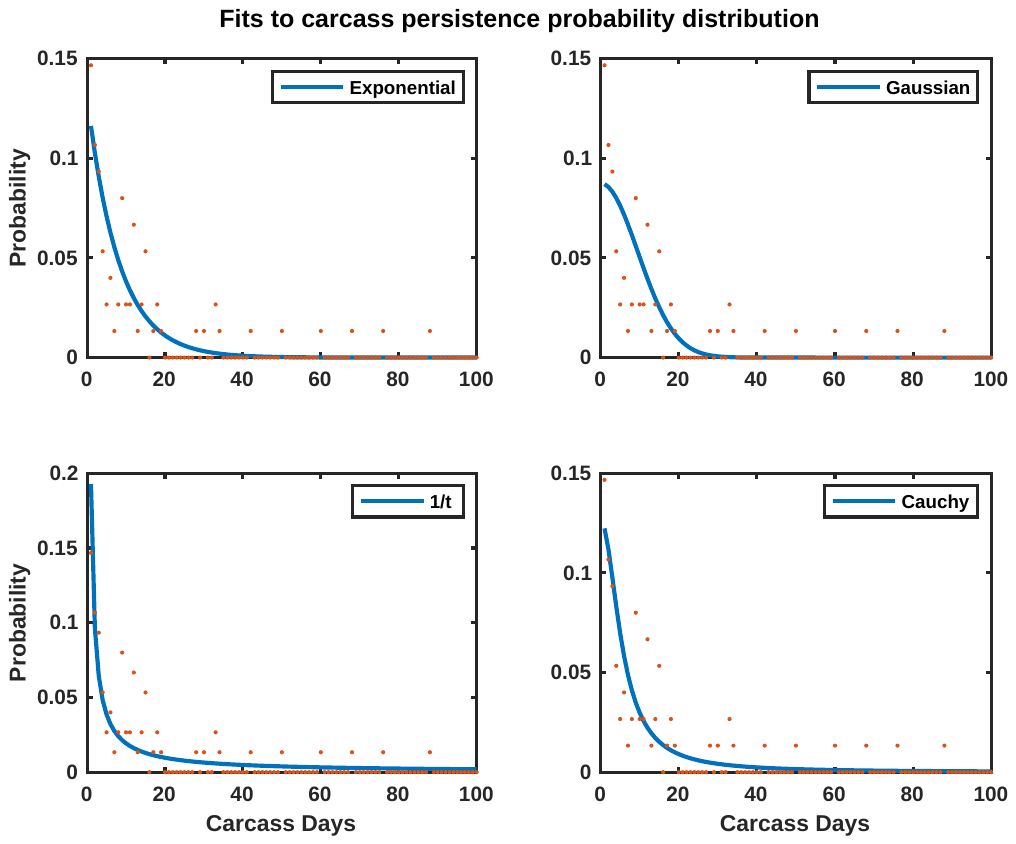

Table S2 Ordinary Least Squares error of fit of various functions for probability of scavenging.

|  |  | Total OLS error |
| --- | --- | --- |
| Exponential distribution | $F\sim e^{-ad}$ | 1.2558 |
| Gaussian distribution | $F\sim e^{-ad^{2}}$ | 1.0000 |
| “1/t” distribution | $F\sim\frac{a}{d}$ | 4.2103 |
| Cauchy distribution | $F\sim\frac{1}{\left( 1+\left( \frac{d}{a} \right)^{2} \right)}$ | 1.1502 |

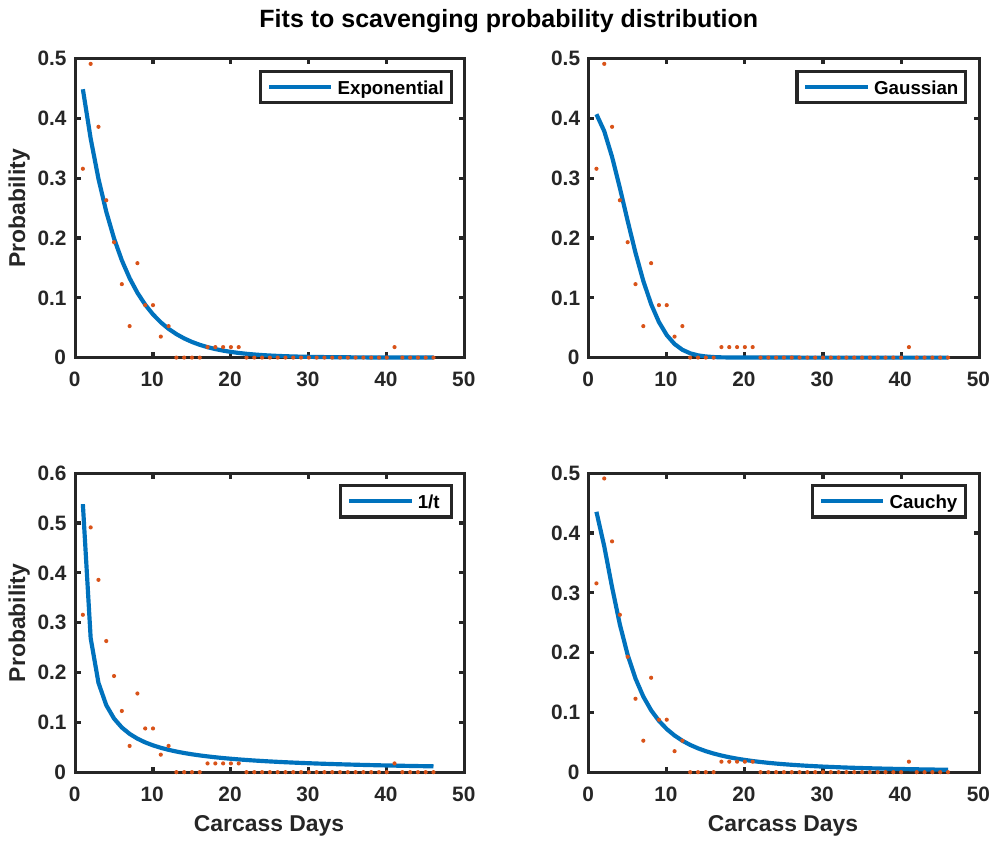

Table S3 Ordinary Least Squares error of fit of various functions for scavenging-hours when scavenging occurs.

|  |  | Total OLS error |
| --- | --- | --- |
| Exponential distribution | $F\sim e^{-ad}$ | 1.0047 |
| Gaussian distribution | $F\sim e^{-ad^{2}}$ | 1.0000 |
| “1/t” distribution | $F\sim\frac{a}{d}$ | 1.1879 |
| Cauchy distribution | $F\sim\frac{1}{\left( 1+\left( \frac{d}{a} \right)^{2} \right)}$ | 1.0014 |

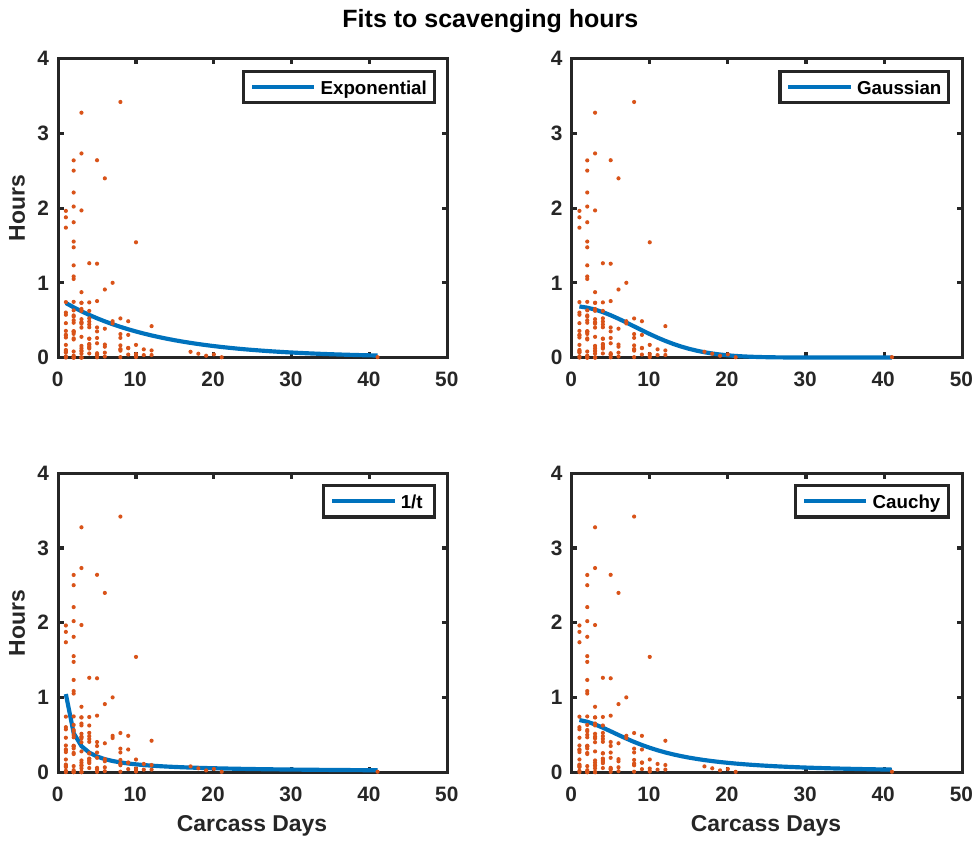

#### Cauchy distribution parameters for final fits

All fits in the manuscript are of the form $F(d)=s\frac{2N}{\pi\gamma\left( 1+\left( \frac{d}{\gamma} \right)^{2} \right)}$ where s is a scale factor, d is the number of days, which can take on positive integer values. N is a normalization factor which assures that the discrete sum over probabilities is 1. As can be seen in tables S2 and S3, the Cauchy distribution does not always have the absolute lowest errors. However, the error differences are very small, and the Gaussian distribution, which does have the lower errors, has the drawback of being effectively zero beyond 3 weeks after which there is some reported activity.

Table S4 Cauchy distribution parameters for final fits.

|  | $\gamma$ | s | Notes |
| --- | --- | --- | --- |
| Carcass persistence probability | 5.5547 | 1 | s=1 because this is a probability distribution with total probability=1 |
| Likelihood of scavenging per carcass-day | 4.3179 | 2.6991 | Here, s corresponds to the average number of days each carcass is scavenged. |
| Number of scavenge-hours | 9.2695 | 8.4854 | Here, s corresponds to the total number of hours the average carcass is scavenged. |
